# Dynamic DnaA-DnaB interactions at *oriC* coordinate the loading and coupled translocation of two DnaB helicases for bidirectional replication

**DOI:** 10.1101/2025.06.05.657979

**Authors:** Takumi Tsuruda, Ryusei Yoshida, Chihiro Hayashi, Kazutoshi Kasho, Shogo Ozaki, Tsutomu Katayama

## Abstract

Bidirectional replication is a conserved principle requiring the coordinated loading and activation of two replicative helicases at the origin. In *Escherichia coli*, the initiator protein DnaA constructs a higher-order initiation complex at the origin *oriC* which locally unwinds the DNA, and recruits DnaB helicase-DnaC loader complexes to the unwound region. We previously demonstrated that the two DnaA subcomplexes formed on *oriC* bind a specific DNA strand of the unwound origin, and tether individual DnaB-DnaC complexes via stable interactions between DnaA domain I and DnaB. A low-affinity DnaA-DnaB interaction mediated by DnaA domain III His136 is essential for DnaB-dependent origin unwinding. Here, we identified DnaB Thr86 as the critical residue mediating this low-affinity interaction. Structural modelling suggests that Thr86 is surface-exposed near the DNA entry site of DnaB. Functional analyses revealed that DnaB Thr86 was specifically required for DnaB loading onto the DnaA-bound strand of the unwound *oriC*. Furthermore, this strand-specific DnaB loading was required for enabling translocation of the opposing DnaB helicase loaded on the DnaA-free strand. Our findings define a novel regulatory mechanism of strand-specific helicase loading, mediated by the low-affinity DnaA–DnaB interactions, which ensures the coordinated loading and translocation of DnaB helicases for bidirectional replication from *oriC*.

## INTRODUCTION

Chromosomal DNA replication initiates bidirectionally at replication origins, a fundamental principle conserved across life (1). Critical steps supporting this principle are the bidirectional loading of two molecules of replicative helicases onto the origin DNA and timely activation of the both of the loaded helicases. A set of replication proteins are recruited to the loaded helicase, constructing a replisome to synthesize daughter strands.

In both bacteria and eukaryotes, the replicative helicases are typically recruited to the origin by specific origin-binding proteins in both bacteria and eukaryotes. In *Escherichia coli*, the initiator protein DnaA specifically binds to the origin DNA, and constructs the initiation complex with a specific DNA bending protein, which induces local DNA unwinding to generate single-stranded DNA (ssDNA) region (2–4). DnaB is then loaded to the ssDNA region through multiple dynamic interactions with the initiation complex (5–11). Similarly, the ORC-Cdc6 complex in budding yeasts and human cells binds to the origin DNA and construct higher-order complexes with the replicative helicase Mcm2-7 (12). Recent studies have reported that yeast Mcm2-7 is loaded onto the double-stranded DNA (dsDNA) through multiple dynamic interactions with the ORC-Cdc6 complex (13,14). Those processes result in loading of the two helicase molecules bidirectionally on the origin DNA. However, the crucial mechanism to ensure timely activation of the both of the loaded helicases remain unclear.

In *E. coli*, the replication origin *oriC* consists of a Duplex Unwinding Element (DUE) and a DnaA-Oligomerization Region (DOR) (Figure 1A) (2–4). The DUE contains three AT-rich 13-mer repeat sequences (L-, M- and R-DUE), and is flanked with a 12-mer AT-cluster region upstream the L-DUE. The upper strand of the ssDUE includes specific sequences TT[A/G]T(T) which bind to DOR-bound DnaA complexes when the DUE is unwound (see below) (11,15,16). The DOR is subdivided into the Left-, Middle-, and Right-DOR (Figure 1A). The Left- and Right-DOR contain clusters of the specific DnaA binding site termed DnaA box, which has 9-mer consensus sequence TTA[T/A]NCACA (17,18). The Left-DOR bears crucial five DnaA boxes termed R1, R5M, τ2, I1 and I2. The Right-DOR also bears five DnaA boxes which are termed R4, C1, I3, C2 and C3. In addition, the interspace between R1 and R5M boxes in the Left-DOR has the IHF binding region (IBR) containing the specific binding sequence of a DNA-bending protein IHF (16). The Left-DnaA subcomplex is constructed by binding of a DnaA pentamer to the five DnaA boxes, which is stimulated by DNA bending caused by IHF binding (Figure 1A and 1D) (7 ,16,19,20).

**Figure 1.**
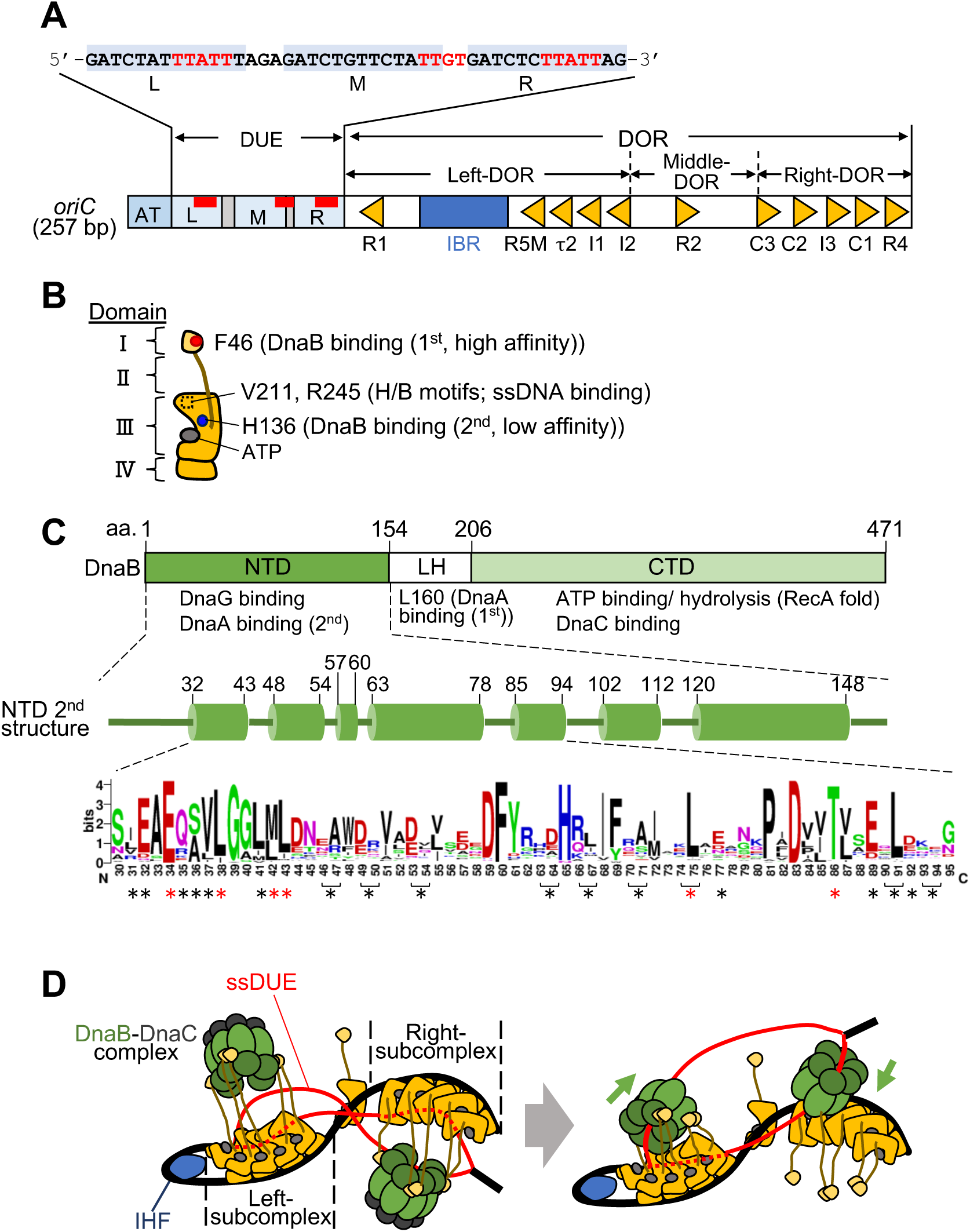
Schematic structures of *oriC*, DnaA and DnaB and the DnaB loading mechanism. (A) Structure of *oriC* (257 bp). *oriC* contains the Duplex-Unwinding Element (DUE; light blue bar) and the DnaA-Oligomerization Region (DOR; open bar). The DUE contains 13-mer repeat sequences (L-, M-, and R-DUE). The sequence of the upper (T-rich) strand of the DUE is shown above the structure. ssDNA binding motifs for DnaA (TT[A/G]T(T)) are highlighted in red. The DOR includes 11 functional DnaA boxes (yellow arrowheads) and an IHF binding region (IBR; dim blue bar). The DOR is divided into Left, Middle, and Right subregions. The AT-cluster upstream the DUE is an auxiliary unwinding region (blue box). (B) A schematic structure of ATP-DnaA. DnaA consists of four domains (I-IV), each indicated by brackets. Domain I tightly binds to DnaB by the F46 residue (red circle). Domain III, an AAA+ domain, is responsible for ATP binding, low-affinity DnaB binding by the H136 residue (blue circle), and the ssDNA binding through the V211 and R245 residues (H/B motifs, dashed squares). (C) Domain structure of DnaB. N-terminal domain (NTD; green), Linker helix (LH; white), and C-terminal domain (CTD; light green) are shown with their specific functions. The middle figure depicts the secondary structure of the NTD, where cylinders represent α-helices. The conservation of amino acid residues within the Ser30-Gly95 region is illustrated using WebLogo. Asterisks represent the amino acid residues substituted with alanine in this study (Table 1). Red asterisks indicate residues that are essential for the complementing activity against the temperature-sensitive *dnaB4*3 mutant cells. (D) Mechanism of DnaB loading by the *oriC*-DnaA complex. The Left-DnaA subcomplex including IHF promotes DUE unwinding and the unwound region is dynamically expanded through interaction between the upper strand of the ssDUE and both the Left- and Right-DnaA subcomplexes (*left panel*). Each DnaA subcomplex binds to a DnaBC complex via a high affinity interaction between DnaA F46 and DnaB L160 residues. Subsequently, DnaBC complexes are loaded into each ssDUE strand, a process thought to require low-affinity DnaA-DnaB interactions (*right panel*). One of the ssDUE-loaded DnaBC complexes localizes to the M/R-DUE region of the lower strand, and the other to the L/M-DUE region of the upper strand. DnaA is depicted as in *panel B*. IHF, DnaB NTD-LH, DnaB CTD and DnaC are depicted respectively by dark blue, green, light green and dark green ovals.

**Table 1.**
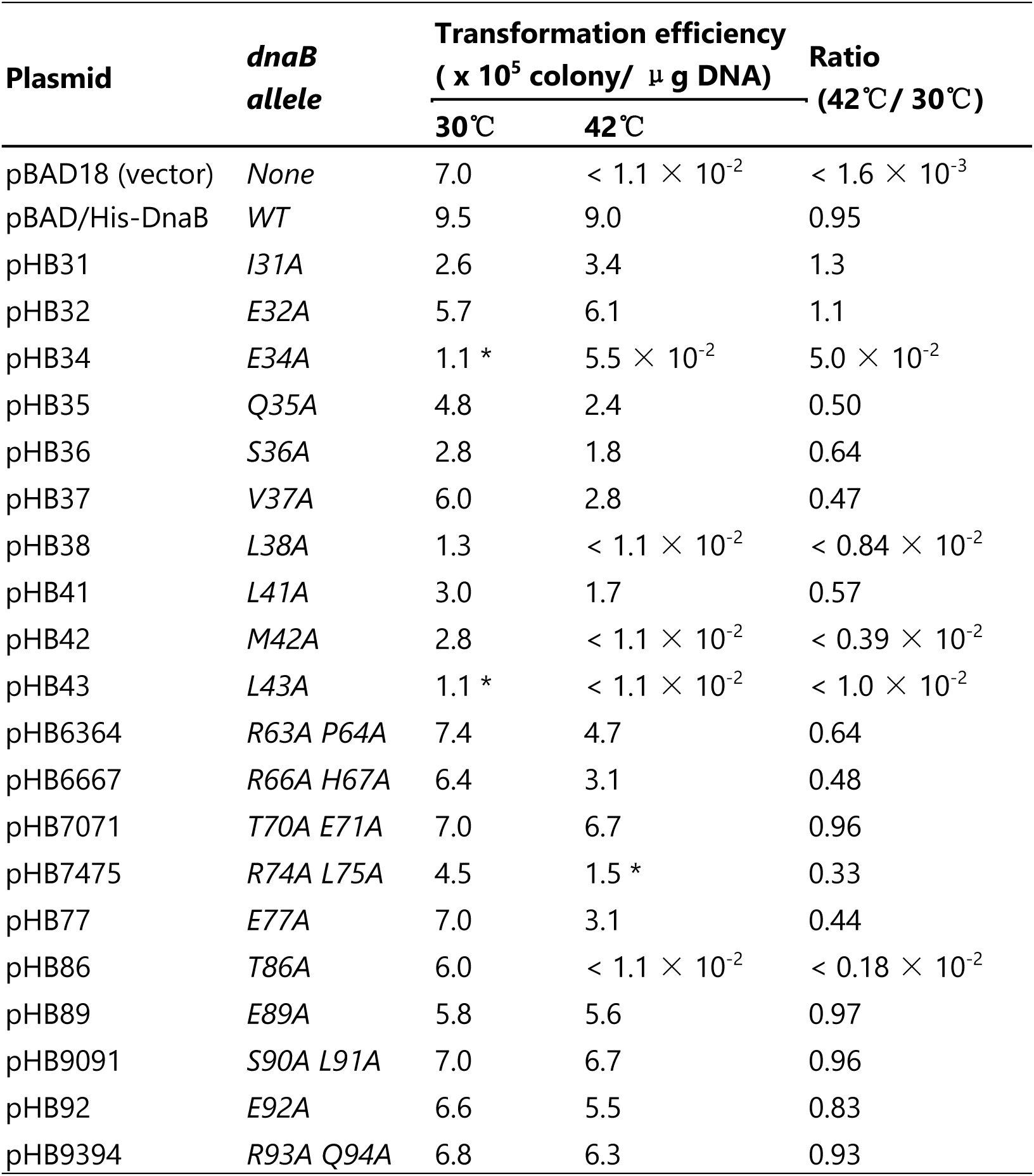
Plasmid complementation test. Plasmid pBAD18 (None) or its pBAD/His-DnaB derivative bearing WT *dnaB* (WT) or the indicated mutant allele was introduced into ME5491 (*dnaB43*) cells. Transformants were incubated at 30°C for 19 hr or 42°C for 14 hr on LB-agar plates containing 0.1% arabinose and 50 mg/ml ampicillin, and then colonies were counted. Transformation efficiency (colony-forming units/μg of DNA) at each temperature, and its ratios (42°C/30°C) are shown. *, only tiny colonies with a diameter of < 0.5 mm were formed.

| Plasmid | *dnaB*<br>allele | Transformation efficiency<br>($\times 10^5$ colony/μg DNA) | | Ratio<br>(42°C/30°C) |
| --- | --- | --- | --- | --- |
|  |  | 30°C | 42°C |  |
| pBAD18 (vector) | *None* | 7.0 | $< 1.1 \times 10^{-2}$ | $< 1.6 \times 10^{-3}$ |
| pBAD/His-DnaB | *WT* | 9.5 | 9.0 | 0.95 |
| pHB31 | *I31A* | 2.6 | 3.4 | 1.3 |
| pHB32 | *E32A* | 5.7 | 6.1 | 1.1 |
| pHB34 | *E34A* | 1.1 * | $5.5 \times 10^{-2}$ | $5.0 \times 10^{-2}$ |
| pHB35 | *Q35A* | 4.8 | 2.4 | 0.50 |
| pHB36 | *S36A* | 2.8 | 1.8 | 0.64 |
| pHB37 | *V37A* | 6.0 | 2.8 | 0.47 |
| pHB38 | *L38A* | 1.3 | $< 1.1 \times 10^{-2}$ | $< 0.84 \times 10^{-2}$ |
| pHB41 | *L41A* | 3.0 | 1.7 | 0.57 |
| pHB42 | *M42A* | 2.8 | $< 1.1 \times 10^{-2}$ | $< 0.39 \times 10^{-2}$ |
| pHB43 | *L43A* | 1.1 * | $< 1.1 \times 10^{-2}$ | $< 1.0 \times 10^{-2}$ |
| pHB6364 | *R63A P64A* | 7.4 | 4.7 | 0.64 |
| pHB6667 | *R66A H67A* | 6.4 | 3.1 | 0.48 |
| pHB7071 | *T70A E71A* | 7.0 | 6.7 | 0.96 |
| pHB7475 | *R74A L75A* | 4.5 | 1.5 * | 0.33 |
| pHB77 | *E77A* | 7.0 | 3.1 | 0.44 |
| pHB86 | *T86A* | 6.0 | $< 1.1 \times 10^{-2}$ | $< 0.18 \times 10^{-2}$ |
| pHB89 | *E89A* | 5.8 | 5.6 | 0.97 |
| pHB9091 | *S90A L91A* | 7.0 | 6.7 | 0.96 |
| pHB92 | *E92A* | 6.6 | 5.5 | 0.83 |
| pHB9394 | *R93A Q94A* | 6.8 | 6.3 | 0.93 |

DnaA consists of four functional domains I-IV (Figure 1B) (21,22). Domain I interacts with several proteins including DnaB, DnaA-assembly stimulator DiaA, and DnaA domain I itself (8,23–26). In particular, the Phe46 residue in this domain is the first and high-affinity interaction site for DnaB (8,26). Domain II is a flexible linker (26). Domain III has AAA+ (ATPase associated with various cellular activities) motif for ATP/ADP binding, ATP hydrolysis, and domain III-domain III interaction (27,28). In addition, the H/B motifs (Val211 and Arg245) bind to the ssDUE TT[A/G]T(T) sequences (7,15,29). Also, the His136 residue is the second and low-affinity interaction site for DnaB (9). Domain IV specifically binds to the dsDNA bearing DnaA box (30).

To initiate the replication, ATP-DnaA molecules and IHF construct a higher-ordered initiation complex on the DOR, including two DnaA pentamers (Left- and Right-DnaA subcomplex) (Figure 1D) (7,11,16,19). The Arg285 residue in DnaA domain III interacts with ATP bound to a flanking DnaA molecule, sustaining cooperative binding in these DnaA pentamers (27,28,31). In the Left-DnaA subcomplex, the DUE is unwound and sharp DNA bending caused by IHF binding at the IBR promotes binding of the H/B motifs in the DnaA pentamer to the ssDUE-upper strand TT[A/G]T(T) sequences, thereby stabilizing the unwound state (11,15,20). This DUE unwinding mechanism is called the ssDUE-recruitment mechanism or DNA loopback mechanism (2–4,7,32). Further extension of the unwound region is stimulated by ssDUE binding of the Right-DnaA subcomplex (Figure 1D) (11).

DnaB helicase is a homohexamer with a ring-like configuration and its protomer consists of three domains (Figure 1C) (33–35). The N-terminal domain (NTD) forms a trimer-of-dimer in the hexamer and interacts with DnaG primase (36–39). The linker helix (LH) region includes the Leu160 residue, which binds to DnaA domain I Phe46, establishing the first and high-affinity interaction between the two proteins (10). The C-terminal domain (CTD) contains a RecA-like ATPase fold (40). Together with LH and CTD, DnaB stably binds to the loader DnaC to construct a DnaB_6_:DnaC_6_ (DnaBC) complex, which causes the conformational change to an open helical configuration, enabling the topological loading of DnaB to ssDNA (34,41–43). Notably, the DnaB NTD is suggested to include a specific site for the second and low-affinity interaction with DnaA His136, which promotes DnaB topological loading to the unwound DUE of *oriC* (44). Following DnaBC loading onto the ssDUE, DnaC is dissociated depending on its interaction with ssDNA, which is stimulated by DnaB-DnaG binding and rNTP (6,34,41,45). Both of the loaded DnaB helicases then translocate in the 5’-to-3’ direction to unwind dsDNA depending on ATP hydrolysis (Figure 1D). This translocation activity is further stimulated by the binding of SSB (Single-Strand-Binding protein) to the expanded ssDNA region (46,47).

DnaB loading onto the ssDUE region within the initiation complex is sequentially promoted by dynamic interactions between DnaA and DnaB. Initially, each DnaA subcomplex binds a DnaBC complex through a high-affinity interaction between DnaA Phe46 and DnaB Leu160 (Figure 1D) (7,8,10,11). On the surface of each DnaB hexamer within the DnaBC complex, three Leu160 residues are exposed, allowing multiple DnaA protomers to interact with a single DnaB hexamer (10). This interaction stabilizes the tethering of the DnaBC complex to the DnaA subcomplexes. Within this dynamic assembly, the flexibility of DnaA domain II permits the DnaBC complexes to move within a certain spatial range (26).

Subsequently, the His136 residues of DnaA domain III in the DnaA subcomplexes are proposed to engage in transient, low-affinity interactions with the DnaBC complexes, guiding them toward the limited ssDNA region of the DUE (9). Notably, only the ssDUE-upper strand, but not the lower strand, is bound by DnaA domain III within the DnaA subcomplexes (Figure 1D) (11,15). For bidirectional replication, two DnaB helicases must be loaded onto these structurally distinct strands independently. To understand the mechanism underlying such strand-specific DnaB loading, the detailed interaction dynamics between the initiation complex and the DnaBC complex must be investigated. However, progress has been hindered by the lack of identification of the specific residues in DnaB that mediate the low-affinity interaction with DnaA.

In the present study, we investigated the DnaB loading mechanism mediated by low-affinity DnaA-DnaB interactions. First, we identified the DnaB residue specifically supporting the low-affinity interaction with DnaA His136 through a combination of *in vivo* mutant screening and *in vitro* functional analyses. The Thr86 residue in the DnaB NTD was found to specifically mediate the low-affinity interaction. Further analyses demonstrated that Thr86 is essential for the second DnaB loading event onto the ssDUE-upper strand, while it plays a non-essential, stimulatory role in the first DnaB loading event on the lower strand. Moreover, using a DnaB T86A mutant and specific *oriC* mutants, we found that the translocation of the first DnaB helicase on the lower strand requires the loading of the second DnaB helicase on the upper strand. These findings reveal a strand-specific DnaB loading mechanism mediated by the low-affinity DnaA-DnaB interactions, a novel regulatory mechanism that ensures bidirectional replication by dynamic coordination for the functions of DnaB helicases.

## MATERIAL AND METHODS

### Bacterial strains, proteins, DNA and plasmids

Strains ME6299 (HfrP4X8) and ME5491 (ME6299 *dnaB43* (Ts)) were described previously (10) and were acquired from the National Bioresource Project (*E. coli*) of the Institute of National Genetics, Japan. N-terminally Hisx6-tagged DnaB was overproduced using pBAD/His-DnaB, Ala-substitution mutations were introduced to the *dnaB* gene in pBAD/His-DnaB by the reverse-direction PCR using mutagenic primers, and the mutations were confirmed by nucleotide sequencing, as we previously described (10). The resultant plasmids are listed in Tables 1 and 2. Purifications of His-DnaB and its derivatives were performed as we previously described (10). The fork DNA was constructed by annealing of synthetic DNA strands (Supplementary Table S1), as we previously described (10). Sequences of synthetic primers used are also shown in Supplementary Table S1. Plasmids bearing *oriC* derivatives were described previously (11) and are listed in Supplementary Table S2.

**Table 2.**
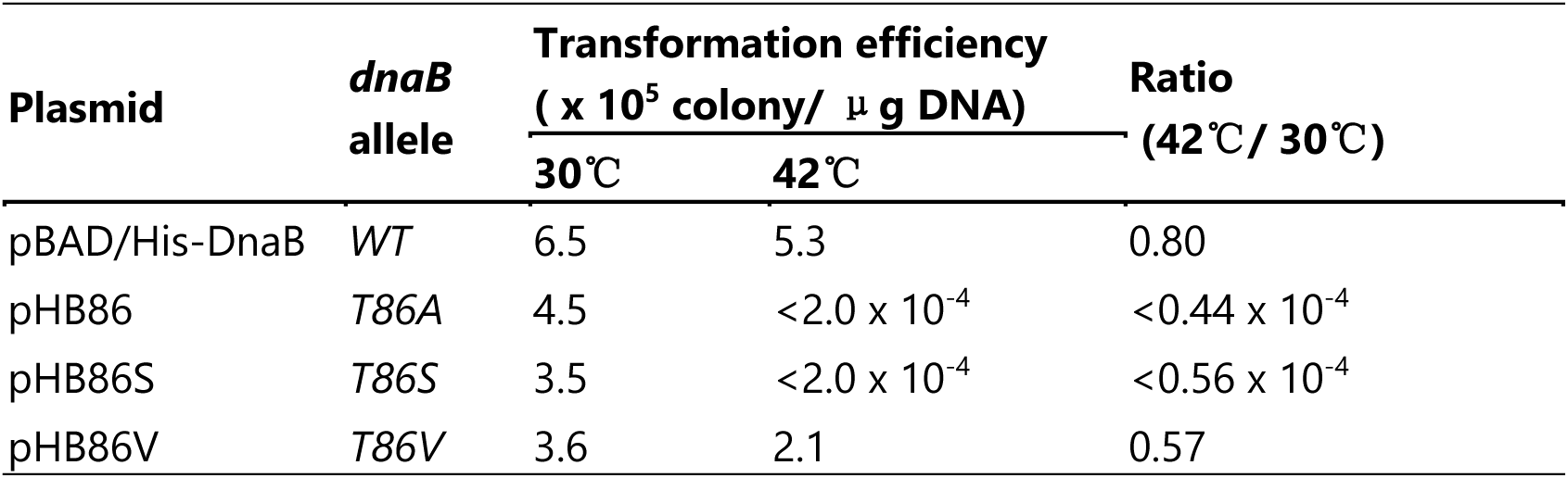
Plasmid complementation test for *dnaB T86* mutants. Experiments were performed as described for Table 1, except for using different plasmids as indicated. Also, different numbers of cells were incubated on agar plates.

| Plasmid | *dnaB*<br>allele | Transformation efficiency<br>($\times 10^5$ colony/μg DNA) | | Ratio<br>(42°C / 30°C) |
| --- | --- | --- | --- | --- |
|  |  | 30°C | 42°C |  |
| pBAD/His-DnaB | WT | 6.5 | 5.3 | 0.80 |
| pHB86 | T86A | 4.5 | $<2.0 \times 10^{-4}$ | $<0.44 \times 10^{-4}$ |
| pHB86S | T86S | 3.5 | $<2.0 \times 10^{-4}$ | $<0.56 \times 10^{-4}$ |
| pHB86V | T86V | 3.6 | 2.1 | 0.57 |

### Form I* assay for DnaB loading

The assay was performed as we previously described, with minor modification of the reaction buffer (5,9,11,43,47). The indicated amounts of His-DnaB and DnaC were incubated for 15 min at 30 °C in Form I* buffer (20 mM Hepes-KOH at pH 7.6, 5 mM magnesium acetate, 5 mM ATP, 125 mM potassium glutamate, 4 mM dithiothreitol, 1 mM EDTA, 10 % glycerol, and 0.04 mg/mL bovine serum albumin) containing 1.6 nM pBS*oriC* or pBS*oriC*ΔDUE, 80 nM DnaA, 42 nM IHF, 76 nM GyrA, 100 nM His-GyrB, and 760 nM SSB. The reaction was stopped by addition of 0.5% SDS, and DNA was purified by phenol–chloroform extraction. Samples were analyzed by 0.65% agarose gel electrophoresis with 0.5xTris-borate-EDTA buffer for 15 h at 23V, followed by ethidium bromide staining.

### Pulldown assays using His-DnaB

DnaB loading onto *oriC* plasmid was assayed as we described previously, with minor modification of reaction buffer (9,10,43). Briefly, 1.6 nM pBS*oriC* or pBS*oriC*ΔDUE was incubated for 9 or 15 min at 30 °C in Form I* buffer containing 80 nM DnaA, 100 nM His- DnaB, 100 nM DnaC, 42 nM IHF and 760 nM SSB, followed by incubation on ice for 15 min with Co^2+^-conjugated magnetic beads (Dynabeads, Thermo Fisher Scientific). Bead-bound materials were collected by magnetic force and washed in buffer containing 100 mM NaCl. Recovered plasmid was eluted in standard SDS sample buffer and analyzed by 1% agarose gel electrophoresis.

The DnaB–DnaC binding assay was performed as we described previously (8,10,43). Briefly, 5 pmol His-DnaB and indicated amount of DnaC were incubated at 4 °C for 15 min in reaction buffer (20 mM Hepes-KOH at pH 7.6, 5 mM magnesium acetate, 1 mM ATP, 90 mM ammonium sulfate, 4 mM dithiothreitol, 1 mM EDTA, 0.1 % Triton X-100, 10 % glycerol, 0.1 mg/mL bovine serum albumin), followed by pulldown with Ni^2+^-conjugated magnetic beads. After washing, recovered proteins were eluted in standard SDS sample buffer and analyzed by SDS-10% polyacrylamide gel electrophoresis (PAGE) and silver staining.

### Helicase activity assay

DnaB helicase activity was assessed using the fork DNA basically as we described previously (10). Briefly, 20 nM fork DNAs were incubated at 37 °C for 40 min in Form I* buffer containing indicated amount of His-DnaB and 100 nM competitor ssDNA. DNA was extracted using phenol/chloroform and analyzed by 6% PAGE.

### Biotin-tagged DOR pull-down assay

The assay was performed as we previously described (7–10). Biotinylated DOR fragment were amplified by PCR, using pBS*oriC* as templates and primers listed in Table S1. 10 nM biotinylated DOR fragment was incubated on ice for 10 min in pull-down buffer (20 mM Hepes–KOH at pH 7.6, 1 mM EDTA, 4 mM dithiothreitol, 5 mM magnesium acetate, 40 mM ammonium sulfate, 20 mM NaCl, 10% glycerol, 0.1% Triton X-100, 1 mM ATP, and 0.1 mg/ml bovine serum albumin), containing 600 nM DnaA, 600 nM His-DnaB, and 600 nM DnaC. Biotinylated DNA-bound materials were recovered using streptavidin-coated beads (Invitrogen), washed in pull-down buffer, dissolved in SDS sample buffer, and analyzed by SDS–11% PAGE and silver staining. In parallel, beads-unbound DNA was extracted by ethanol precipitation and subjected to electrophoresis for quantitative control of beads-bound DNA

### Gel filtration analysis using spin-column

For the DnaB-loading reaction, 40 nM His-DnaB was incubated for 10 min at 30 °C in buffer P (60 mM Hepes-KOH at pH 7.6, 8 mM magnesium acetate, 0.1 mM zinc acetate, 125 mM potassium glutamate, 5 mM ATP, 30 % glycerol) containing 1.6 nM pBS*oriC* or pBS*oriC*ΔDUE, 40 nM DnaA, 100 nM DnaC and 16 nM IHF, followed by incubation for 3 min at 4 °C in the presence of 500 mM NaCl. The reaction was applied into a Sephacryl S-400 HR (Sigma-Aldrich) equilibrated with buffer P containing 500 mM NaCl and 0.05 % Triton X-100, and the plasmid DNA and its bound proteins were isolated into the void by a brief centrifugation (3000 rpm, 3 min) at 4 °C. The DNA and proteins were analyzed by 1 % agarose gel electrophoresis, and 10 % SDS-PAGE, respectively.

### P1 nuclease footprint

The indicated amounts of DnaB K236A ATPase-defective mutant or WT DnaB were incubated for 9 min at 30 °C in buffer P containing 1.6 nM pBS*oriC*, 40 nM DnaA, 100 nM DnaC, 16 nM IHF and 0.32 mg/mL bovine serum albumin, followed by further incubation for 5 min with 100 units P1 nuclease (New England Biolabs). The cleaved positions of plasmid DNA were analyzed by primer extension reaction using 5’-^32^P-labeled primers listed in Table S1, 6% sequencing-gel electrophoresis, and a Typhoon FLA9500 image analyzer (GE Healthcare). The position of each DUE element (L, M, and R) and the AT-cluster region was determined using Sanger sequence with the identical primer.

## RESULTS

### Determination of functional residues in DnaB N-terminal domain

Previous studies have suggested that the DnaB Met1–Asp156 fragment interacts with the DnaA Lys135–Gln148 fragment (44) and that DnaA His136 plays a crucial role in the low-affinity interaction with DnaB (9). Based on these, we presumed that a specific residue within the DnaB Met1–Asp156 region mediate the low-affinity interaction. To identify functionally important residues within this domain, we performed a plasmid complementation test using *dnaB43* temperature-sensitive mutant cells. These cells were transformed with plasmid encoding wild-type (WT) DnaB or DnaB variants carrying alanine substitutions at candidate residues predicted to be surface-exposed based on structural data from various DnaBC complexes (38,48–50) (Figure 1C).

While *dnaB43* cells bearing the vector (pBAD18) preserved the defect in colony formation at a restrictive temperature (42°C), *dnaB43* cells bearing plasmid with WT *dnaB* fully restored it under these conditions (Table 1), consistent with our previous results (10). Notably, plasmids carrying *dnaB* alleles with E34A, L38A, M42A, L43A, R74A-L75A, or T86A substitutions failed to fully complement the temperature sensitivity of *dnaB43* cells (Table 1, Figure 1C).

A cryo-EM structure of the DnaBC complex (PDB: 6QEL) suggests that the side-chains of Leu43, Arg74, Leu75, and Thr86 residues are surface-exposed whereas those of Glu34, Lue38, and Met42 are largely buried (34) (Supplementary Figure S1). In addition, the DnaB Arg74 is indicated to be important for helicase activity by eliciting the single-stranded DNA to the exterior of DnaB (51). Therefore, we focused on the other three residues, Leu43, Leu75, and Thr86, for further analysis.

### The DnaB Thr86 residue is essential for the replication initiation

To test the importance of the three DnaB residues for replication initiation, we assessed the activity of DnaB mutant proteins using the form I* assay. In this *in vitro* assay, DnaB is loaded onto *oriC*-region of supercoiled *oriC* plasmid (form I) by the DnaA-IHF-*oriC* complex. Depending on DnaB translocation expanding ssDNA region, DNA gyrase turns the *oriC* plasmid into its topoisomer (form I*) (Figure 2). This topoisomer migrates at a rate different from the form I plasmid in agarose gel electrophoresis (5,9,11,43,47). Consistently, WT DnaB was active in the form I* formation of *oriC* plasmid in a DnaA-dependent manner (Figure 2BC). By contrast, DnaB T86A protein was severely impaired in the form I* formation (Figure 2BC), suggesting that the Thr86 residue plays a crucial role in the DnaB helicase functions at *oriC*. DnaB L43A protein exhibited about a half the activity of WT, and DnaB L75A protein showed a similar activity to WT.

**Figure 2.**
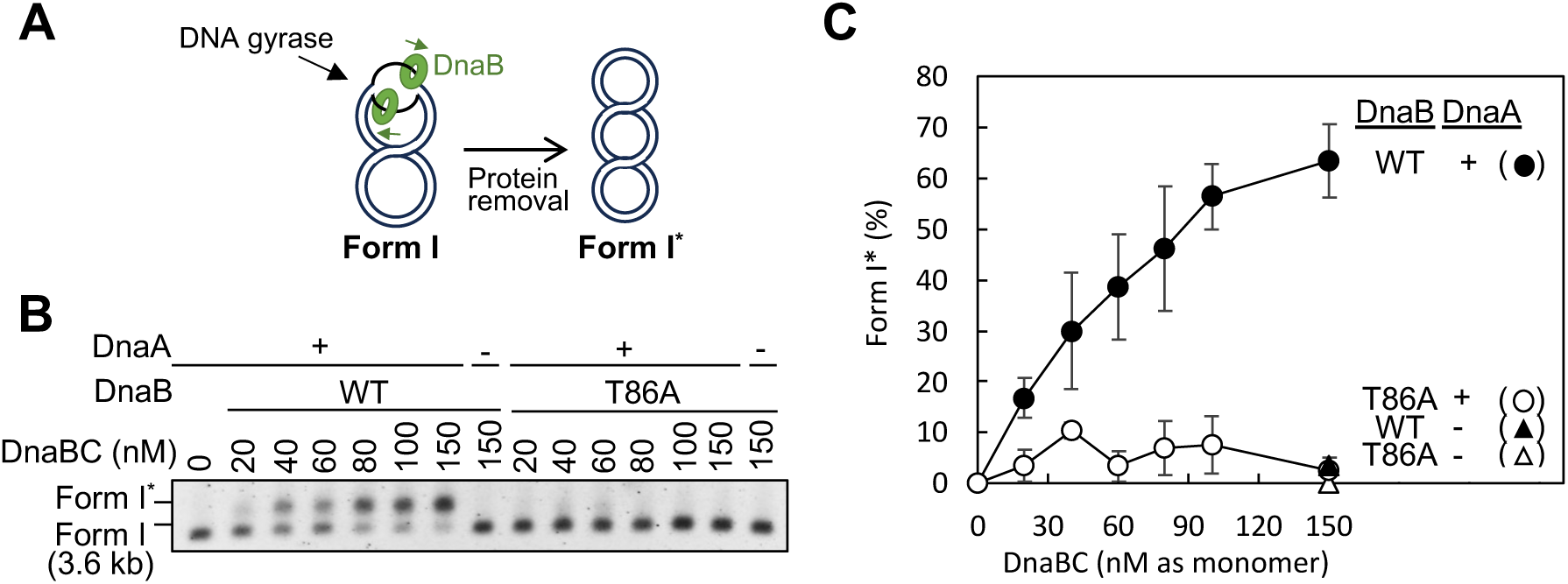
Replication initiation and DnaB loading activity of DnaB T86A mutant. (A) Principle of Form I* assay for detecting the replication initiation (B and C). DnaB helicases are loaded onto the *oriC*-containing plasmid (Form I) in a DnaA-dependent manner. Subsequent translocation of the helicases along the DNA in concert with DNA gyrase generates additional negative supercoiling, converting the plasmid to a more negatively supercoiled form (Form I*). (B and C) Indicated amounts of His-DnaB and DnaC (DnaBC) were incubated for 15 min at 30 °C in the presence of ATP-DnaA, IHF, SSB, and gyrase with 1.6 nM *oriC* plasmid pBSoriC. Following incubation, the plasmid was purified and analyzed by 0.65% agarose gel electrophoresis. A representative gel image from three independent experiments is shown in the black/white-inverted mode (B). Migration positions of Form I and Form I* are indicated. Band intensities were quantified by densitometric scanning and the percentages of Form I* relative to total input DNA are shown as ‘Form I* (%)’ (C). Means and standard deviations (SDs) are shown (n = 3).

To elucidate the functional specificity of DnaB Thr86, we assessed the helicase activity using a fork DNA substrate which consisted of 30-nt ssDNA tails conjugated to a 49-bp dsDNA fragment, as we previously performed (10). DnaB T86A protein unwound this fork DNA at the comparable level to that of WT DnaB (Figure 3AB). In addition, we analyzed the DnaC binding activity by the pulldown assay using affinity recovery of His-tagged DnaB (His- DnaB) as we previously performed (10). WT His-DnaB and its T86A derivative exhibited the similar DnaC-binding activity (Figure 3CD). Taken these results together, we concluded that the DnaB T86A protein is impaired specifically in DnaB helicase-loading to *oriC* and hypothesized that DnaB Thr86 is crucial for the low-affinity DnaA-DnaB interaction. As shown in Figure 2, the severe inhibition of DnaB loading due to T86A is consistent with our previous result demonstration the essential role of DnaA His136 in form I* formation (9).

**Figure 3.**
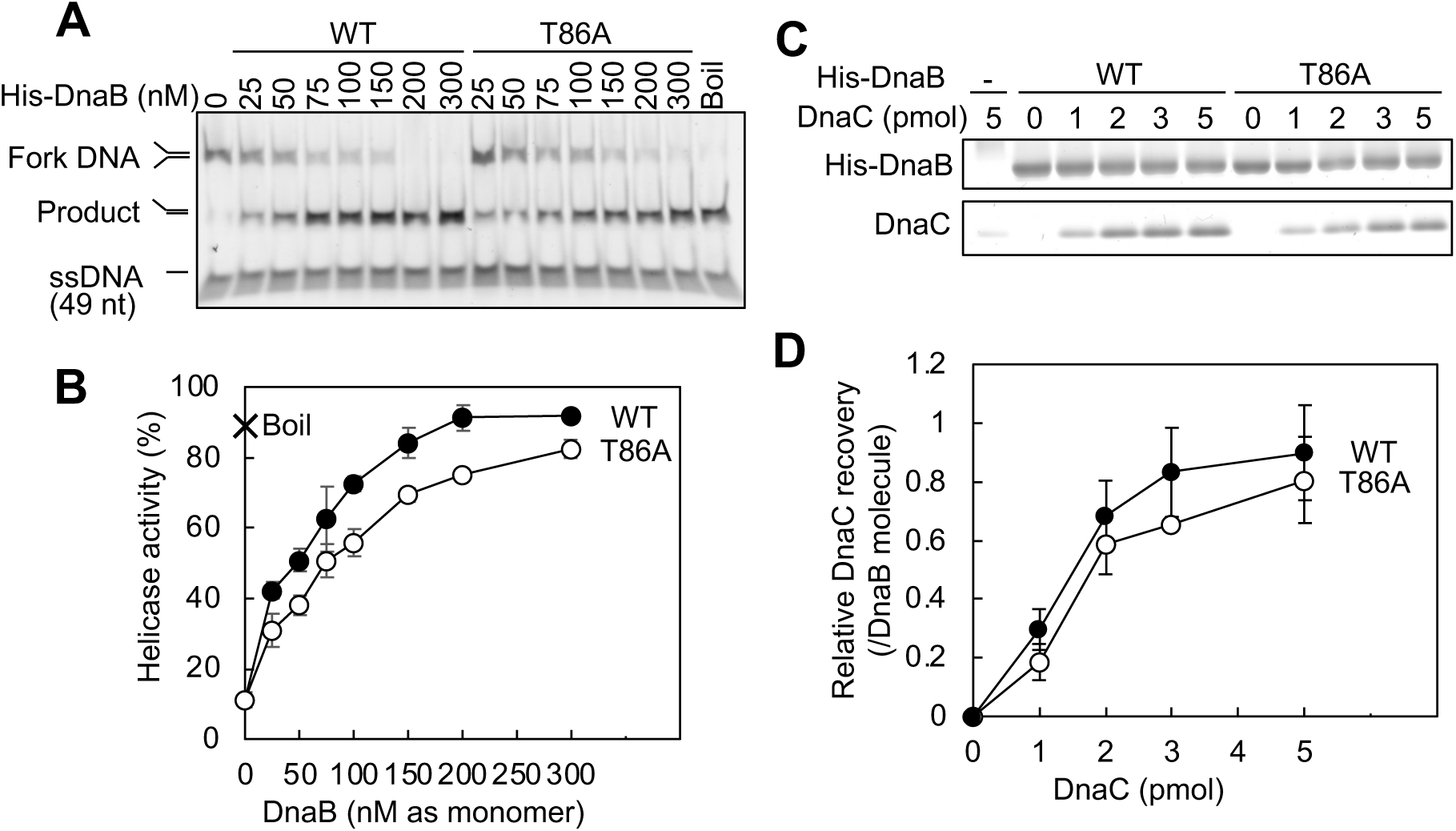
Activities of DnaB T86A in helicase action and DnaC binding. (A and B) Helicase assay of DnaB T86A. Indicated amounts of His-DnaB were incubated with 20 nM the fork DNA substrate for 40 min at 37 °C in the presence of 100 nM competitor ssDNA, which anneals with one of the unwound fork DNA strands. Following incubation, DNA was purified and analyzed by PAGE. For the fully unwound control, fork DNA and competitor ssDNA were incubated for 3 min at 95 °C (Boil), followed by PAGE. A representative gel image is shown in the black/white inverted mode (A). Band intensities were quantified by densitometric scanning, and percentages of the unwound product relative to total input fork DNA are shown as ‘Helicase activity (%)’ (B). Means and standard deviations (SDs) are shown (n = 3). (C and D) His-DnaB pulldown assay for DnaC binding of DnaB T86A. His-DnaB (5 pmol) was incubated with the indicated amounts of DnaC on ice for 10 min. His-DnaB was pull-downed using Co2+-conjugated magnetic beads, and the recovered proteins were analyzed by SDS-PAGE and silver staining. Representative gel images of DnaB and DnaC recoveries from three independent experiments are shown (C). The numbers of recovered DnaB and DnaC molecules were deduced using quantitative standards, and molar ratios of DnaC to DnaB monomer were calculated and shown as ‘Relative DnaC recovery (/DnaB molecule)’(D). Means and standard deviations (SDs) are shown (n = 3). Abbreviations: -, no addiction of His-DnaB

To elucidate the functional-structure relationship in DnaB Thr86, we constructed DnaB T86S and DnaB T86V proteins for analysis by the complementation assay (Table2). The serine substitution, which retains a hydroxyl group similar to threonine, failed to complement the *dnaB43* function at 42°, suggesting that the importance of the Thr86 residue does not stem from its hydroxyl group. In contrast, the valine substitution, which possesses a methyl group, retained the complementation ability. These results are consistent with the idea that the methyl group of the Thr86 residue is crucial for the low-affinity DnaA-DnaB interaction.

### DnaB Thr86 residue is responsible for the low-affinity DnaA-DnaB interaction

DnaB loading to *oriC* ssDNA requires both high-affinity and low-affinity DnaA-DnaB interactions. First, DnaB is recruited to the DnaA-*oriC* complex by the high-affinity interaction between the DnaB LH region Leu160 site and the DnaA domain I Phe46 site (8,10) (Figure 1). Next, DnaB is loaded onto *oriC* ssDNA through the low-affinity interaction with the DnaA domain III His136 site (9) (Figure 1). Therefore, we tested whether the DnaB Thr86 residue contributes to these high-affinity and low-affinity interactions using purified proteins.

First, we analyzed the high-affinity interaction of DnaB T86A using *oriC*-pulldown assay as we previously performed (7,8,10). In the presence of DnaA, DnaB molecules are co-recovered with the biotin-tagged *oriC* DOR fragment (bio-DOR) depending on construction of the DnaA-DOR complexes (Figure 4A). Note that the majority of DnaB recovery in this assay is attributed to the high-affinity DnaA -DnaB interaction (8,10). In the presence of bio- DOR and DnaA, DnaB was recovered at a basal level, and DnaC enhanced the DnaB recovery (Figure 4B), fundamentally consistent with our previous results (8,10). DnaC binding changes the DnaB hexamer structure, which likely enhances the high-affinity DnaA-DnaB interaction. In both WT DnaB and DnaB T86A, approximately 10 molecules of DnaA were recovered per DOR, suggesting that the DnaB T86A mutation has substantially no effect on the formation of the DnaA complex on DOR (Figure 4BC). Furthermore, the recovery of DnaB T86A was 10– 14 molecules as monomers per *oriC*, corresponding to about two hexamers with a consideration to minor loss during purification. This is comparable to the recovery of DnaB WT, indicating that DnaB Thr86 is not substantially involved in the high-affinity interaction with DnaA. In addition, these results are consistent with our previous results indicating that two DnaBC-hexamer complexes tightly bind to the single *oriC*-DnaA complex (7) (Figure 1D).

**Figure 4.**
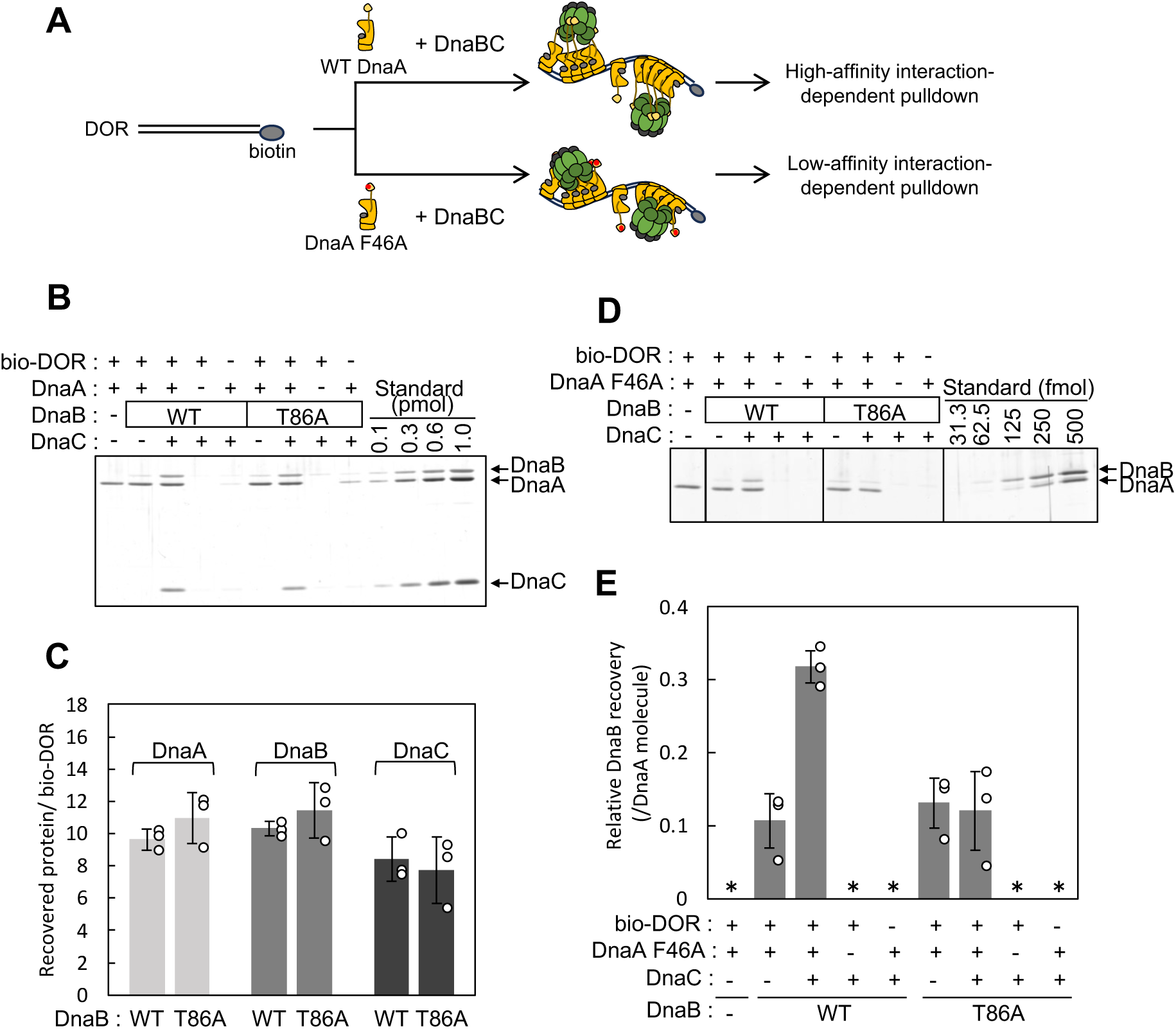
DnaA-DnaB interaction analysis by DOR pulldown. (A) Principle of the bio-DOR pulldown assay. The biotin-tagged DOR (bio-DOR) was incubated with WT DnaA or DnaA F46A in the presence of DnaBC complex, followed by pulldown using streptavidin beads and analysis using SDS-PAGE and silver staining. The bio-DOR complexes with WT DnaA bind the DnaBC complexes with high affinity while those with DnaA F46A and the DnaBC complexes represent the low affinity interaction. Biotin was conjugated to the right end of the full-length DOR fragment (Figure 1). (B and C) Bio-DOR pulldown assay for analyzing the high-affinity DnaA-DnaB interaction. The bio-DOR fragment (10 nM) was incubated on ice for 10 min with (+) or without (-) 600 nM each of DnaA, DnaC and His-DnaB or His-DnaB T86A, followed by pull-down assay as described above. A representative gel image from three independent experiments is shown (B). The recovered protein amounts were deduced using quantitative standards and ratios of each protein to the recovered bio-DOR are shown (C). Data points, means and standard deviations (SDs) are also shown (n = 3). (D and E) Bio-DOR pulldown assay for the low-affinity DnaA-DnaB interaction. This assay was similarly performed using DnaA F46A in place of WT DnaA. A representative gel image from three independent experiments is shown (D). The recovered protein amounts were deduced using quantitative standards and ratios of DnaB to DnaA are shown (E). Data points, means and standard deviations (SDs) are shown (n = 3). *, not detected.

Next, to analyze the low-affinity interaction, we used DnaA mutant proteins bearing F46A which specifically exclude the effect of the high-affinity interaction (Figure 4A). In this experimental setup, the DnaB recovery should depend solely on the low-affinity interaction. A minimum level of recovery of WT DnaB was observed in the absence of DnaC, and DnaC enhanced DnaB recovery by approximately threefold (Figure 4DE). These results suggest that the low-affinity DnaA-DnaB interaction also is stimulated in a DnaC-dependent manner, likely due to a conformational change in DnaB induced by DnaC binding (43). In contrast, DnaB T86A was recovered only at a minimum level regardless of the presence of DnaC (Figure 4DE), suggesting the idea that the DnaB Thr86 residue supports the functional low-affinity interaction depending on DnaC binding. This idea is consistent with the structural characteristics that DnaBC complex takes on an open helical structure and DnaB Thr86 residues are exposed on the surface near the ring-opening site (34,52) (Supplementary Figure S2) (see Discussion). The minimum level recovery observed without DnaC would be due to non-specific interactions and would not be related to the functional DnaA-DnaB interactions for the DnaB loading to *oriC*.

### The DnaB Thr86 residue is crucial for the second DnaB loading onto *oriC* ssDNA

To clarify the specific role for DnaB Thr86 in the low-affinity interaction, we evaluated the topological loading activity onto *oriC* ssDNA using the His-DnaB pulldown assay. When His-DnaB is topologically loaded onto *oriC* plasmid by encircling the ssDNA, the His-DnaB- loaded *oriC* plasmid can be co-recovered by the His-tag pulldown assay under the high-salts conditions (9,10,43) (Figure 5A). In the presence of WT His-DnaB, WT *oriC* plasmid was co-recovered depending on DnaA and DnaC, whereas the *oriC* ΔDUE plasmid showed little recovery even in the presence of DnaA and DnaBC, supporting the idea that this assay specifically detects the topological loading of His-DnaB onto *oriC* ssDNA (Figure 5BC). At a reaction time of 9 minutes, DnaB T86A exhibited approximately half loading activity of WT (Figure 5B), suggesting the specific importance of the DnaB Thr86 residue for the DnaB topological loading. At a reaction time of 15 minutes, DnaB T86A displayed the loading activity at the comparable level to that of WT DnaB (Figure 5C). In this pulldown assay, at least one DnaB hexamer loading event per *oriC* plasmid should be sufficient for the co-recovery of the plasmid. Considering these results, the intact helicase activity of DnaB T86A, and its compromised activity in the form I* assay (Figure 2 and 3), it is possible that the DnaB T86A mutant forms an abortive complex on ssDUE by the topological loading of only a single DnaB hexamer on the ssDUE region. We thus infer the hypothesis that the first loaded DnaB hexamer can not fully translocate on DNA as a helicase unless the second DnaB hexamer is loaded on ssDUE region.

**Figure 5.**
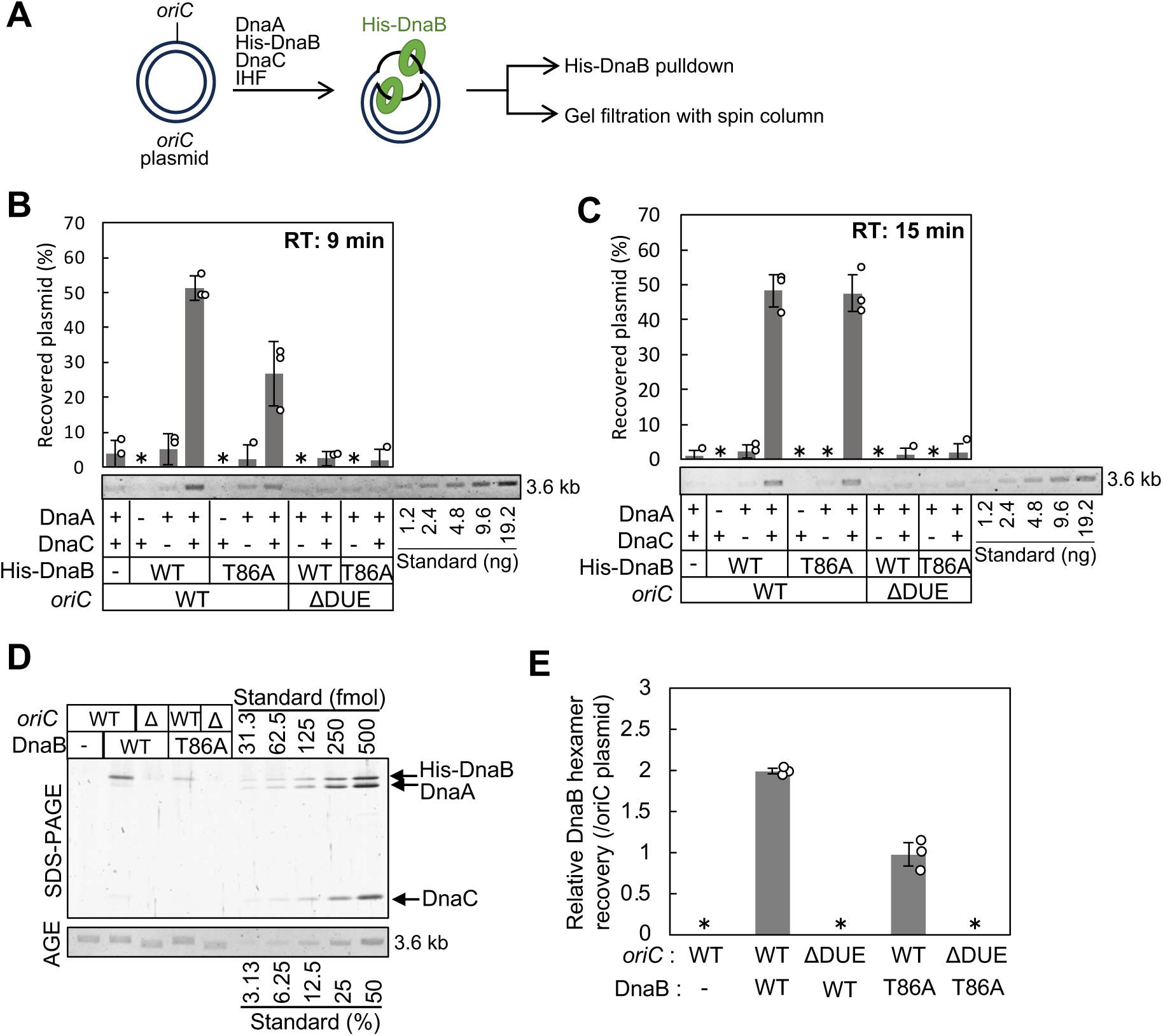
DnaB-loading activity to *oriC* ssDUE. (A) Principle of DnaB-loading assay using His-DnaB pulldown and gel-filtration. The *oriC* plasmid pBSoriC was incubated with the indicated proteins to allow His-DnaB loading onto ssDUE. The His-DnaB-loaded plasmid was isolated by Co^2+^-conjugated beads in high-salts buffer. Alternatively, it was isolated using gel-filtration spin column in high-salts buffer. In these assays, DNA gyrase was absent, thereby restricting translocation of the loaded DnaB. (B and C) His-DnaB pulldown assay for DnaB loading activity. pBSoriC (1.6 nM) was incubated for 9 min (B) or 15 min (C) at 30 °C with IHF and SSB in the presence of ATP-DnaA (80 nM), His-DnaB (100 nM), and DnaC (100nM). His-DnaB-bound plasmid was recovered using Co^2+^-conjugated magnetic beads, washed with buffer containing 100 mM NaCl and eluted in sample buffer. The *oriC* plasmid co-recovered with His-DnaB was analyzed with quantitative standards by agarose gel electrophoresis. Three independent experiments were performed for each incubation time. Representative gel images are shown in the black/white inverted mode. The percentages of recovered DNA relative to the total input are shown as ‘Recovered plasmid (%)’. Data points, means and standard deviations (SDs) are also shown (n=3). When the recovery was below the detectable level, the data points were not shown and treated as zero in the calculation of mean values. Abbreviations: -, no addition of proteins; +, addition of proteins; *, not detected; RT, reaction time. (D and E) Gel-filtration assay for DnaB loading activity. pBSoriC (1.6 nM) was incubated for 15 min at 30 °C in the presence of ATP-DnaA (40 nM), His-DnaB (40 nM), DnaC (100 nM) and IHF (16 nM). pBSoriC and bound proteins were isolated by gel-filtration spin column in buffer containing 500 mM NaCl. Portions of eluted samples were analyzed by agarose gel electrophoresis (AGE) and ethidium bromide staining for DNA and by SDS-PAGE and silver staining for proteins. Representative gel images from three independent experiments is shown (C). The recovered protein and DNA amounts were deduced using quantitative standards and the ratios of DnaB hexamer to pBSoriC molecules are shown (D). Data points, means and standard deviations (SDs) are also shown (n = 3). *, not detected.

To elucidate the above hypothesis, we quantified the stoichiometry of loaded DnaB hexamers per *oriC*. Gel filtration of reactions including *oriC* plasmid, IHF, DnaA, and DnaBC complex allows the separation of *oriC* plasmid loaded DnaB hexamers from DNA-free DnaB hexamers (Figure 5A). For WT DnaB, approximately two hexamers per plasmid were recovered in a DUE-dependent manner (Figure 5DE), consistent with previous results (6,11). In contrast, for DnaB T86A, only about one hexamer per plasmid was recovered even after incubation for 15 min (Figure 5DE), suggesting that the DnaB Thr86 residue is crucial for the loading of the second DnaB hexamer onto *oriC.* These results well coincide with the hypothesis described above.

### The DnaB Thr86 residue is required for the DnaB loading onto the upper ssDUE strand

To determine the strand-specificity of the second DnaB loading inhibited by the DnaB T86A mutation, we performed footprint analysis using nuclease P1. In this assay, ssDNA-specific nuclease P1 cleaves the unwound DUE region of *oriC* plasmid, which can be detected by primer extension, and DnaB loading site is identified by its protection from cleavage (Figure 6A). To fix DnaB at the loaded site, we used DnaB K236A mutation which abolished ATPase activity and thus translocation while preserving its ssDNA-loading activity (6,11). In the presence of IHF, DnaA, DnaB K236A protein and DnaC, protection was predominantly observed in the R-side M region of the ssDUE-lower strand, while cleavage at the AT-cluster—L-DUE boundary of the upper strand was enhanced (Figure 6BC), consistent with previous KMnO_4_ modification data (6,11,53). This enhancement in cleavage is likely caused by stabilization of the unwound state in this region due to physical collision by DnaB loaded on both strands of ssDUE.

**Figure 6:**
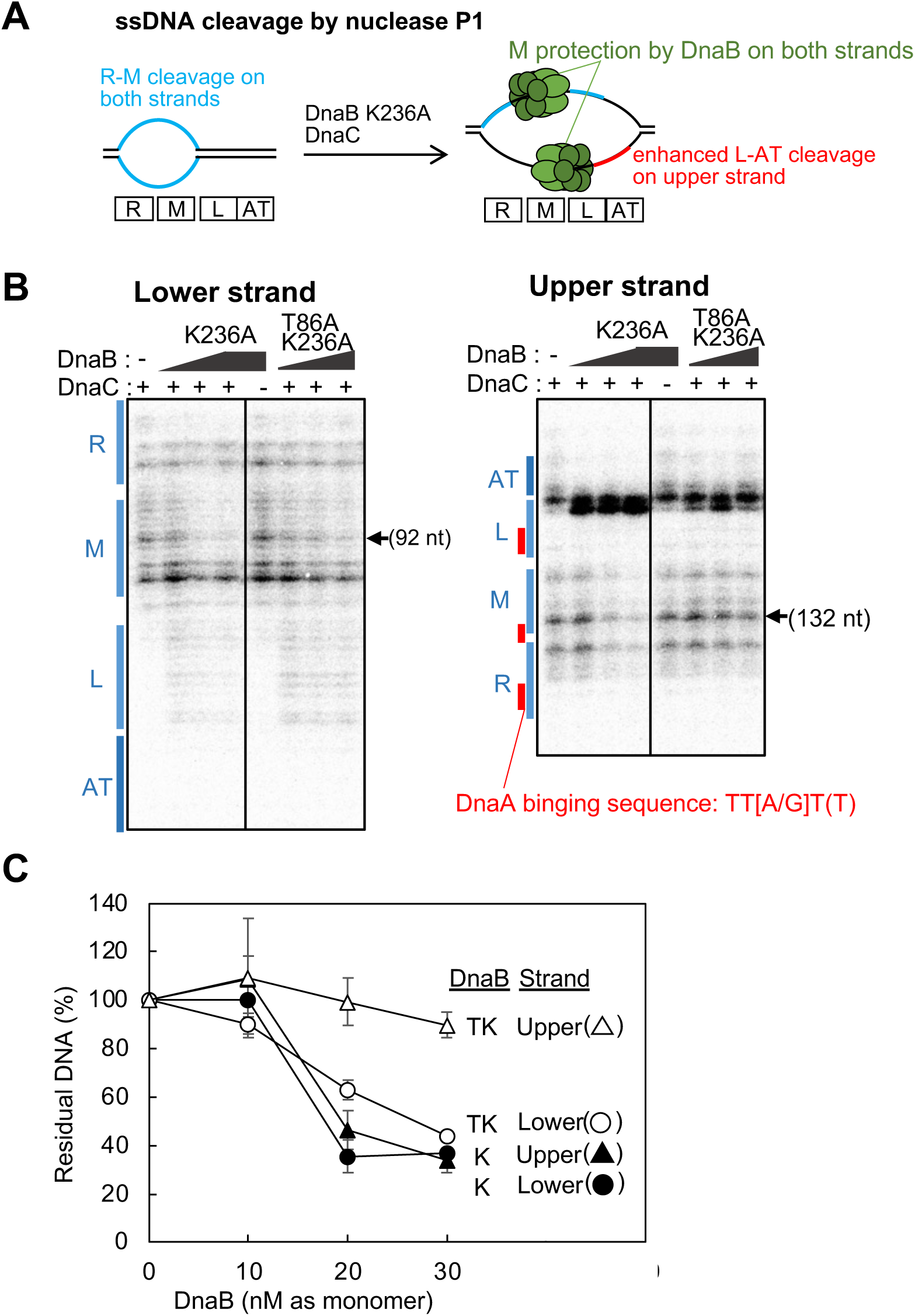
Strand specific analysis of DnaB loading by P1 nuclease footprint experiments. (A) Principle of P1 nuclease footprint experiment. The unwound DUE region in the DnaA-IHF-*oriC* complexes is susceptible to cleavage by P1 nuclease. DnaB K236A ATPase mutant can be loaded onto ssDNA but remains fixed at the initial loading site without translocation. Upon loading, DnaB protects the ssDUE region from the cleavage. As a result, this protection occurred predominantly for M-DUE of the both strands. (B and C) pBSoriC (1.6 nM) was incubated at 30°C for 9 min with IHF (16 nM), ATP-DnaA (40 nM) and 0-30 nM of His-DnaB K236A or His-DnaB T86A K236 in the presence (+) or absence (-) of DnaC (100 nM), followed by incubation with P1 nuclease for 5 min. The resulting DNA was analyzed by primer extension experiments and the products were resolved on 6% sequencing gels (B). Positions of each ssDUE subregion (L, M, and R)and the AT-cluster (AT) are indicated by blue lines at the left side of each gel. Band intensities of specific cleavage products (92 nt for the lower strand and 132 nt for the upper strand), which are indicated by allows, were quantitatively analyzed. The percentages of band intensities relative to those in the absence of DnaB are shown as “Residual DNA (%)” (C). Data points, means and standard deviations (SDs) are also shown (n = 3). Abbreviations: K, DnaB K236A; TK, DnaB T86A K236A

Notably, although the previous KMnO_4_ modification experiments failed to determine the initial loading site of DnaB on the upper strand due to the specificity of its nucleotide sequence (6,11,53), the present footprint analysis demonstrated that the M region of the ssDUE-upper strand was predominantly protected in a DnaB K236A-dependent manner (Figure 6BC). It should be noted that a large part of this protected region lacks the DnaA-binding TT[A/G]T(T) sequences. As such, DnaB K236A with intact Thr86 protected the M-DUE regions of both strands. In contrast, the DnaB T86A K236A double mutant protein failed to protect of the upper-strand M-DUE region, while still fully protecting the lower-strand M- DUE region (Figure 6BC). In the upper strand, the enhancement of cleavage in the AT-cluster—L-DUE boundary was also reduced. These results suggest that the DnaB T86A K236A protein is loaded substantially only to the lower-strand M-DUE region and support the idea that the low-affinity interaction mediated by DnaB Thr86 is specifically crucial for DnaB loading onto the ssDUE-upper strand. Under these experimental conditions of P1 nuclease concentration, short DNA fragments corresponding to the AT-cluster of the lower strand and the late-half portion of R-DUE region of the upper strand were not detected.

### DnaB T86A protein loaded onto the lower strand is unable to translocate

To assess the translocation of DnaB loaded onto the ssDUE-lower strand, we performed the P1 nuclease footprint analysis using the WT DnaB and its T86A derivative. If DnaB helicase is loaded onto the lower strand of DUE and translocates along the DNA with helicase activity, the unwound region should be expanded toward the AT-cluster region. As expected, in the presence of WT DnaB, but not DnaB K236A, the level of the cleavage in the AT-cluster region was increased, suggesting translocation of WT DnaB along the lower strand (Figure 7AB). In this assay, the absence DNA gyrase likely restricted the translocation of DnaB beyond the AT-cluster region. Notably, compared to WT DnaB, DnaB T86A mutation reduced the enhanced cleavage in the AT region (Figure 7AB), suggesting that the translocation of DnaB T86A loaded on the lower strand was inhibited. Given the similar helicase activities between WT DnaB and DnaB T86A proteins (Figure 3AB), the results support the idea that the loading of the second DnaB helicase to the upper strand is required for relieving the inhibition of translocation of the first DnaB helicase loaded on the lower strand.

**Figure 7.**
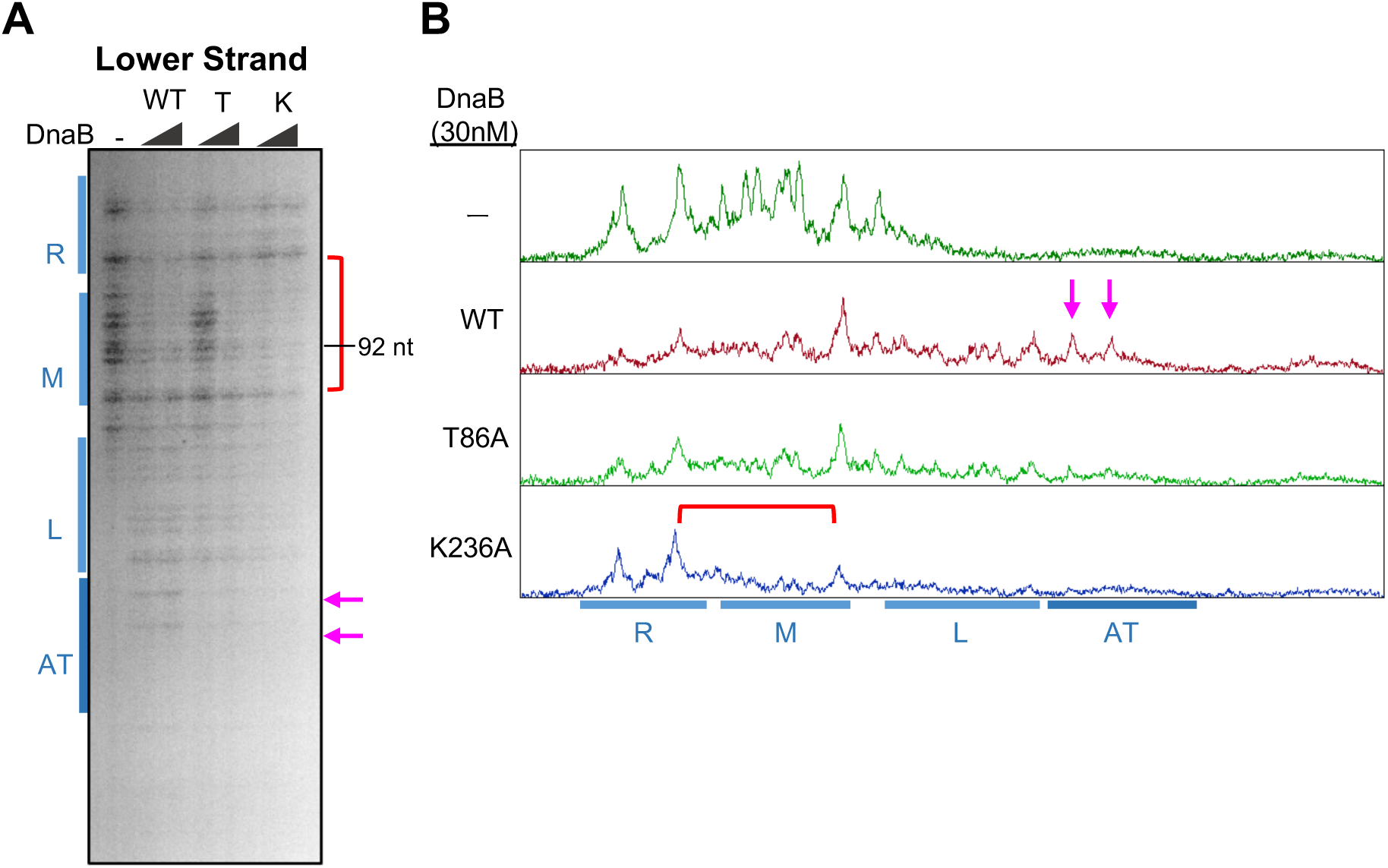
Analysis of the unwinding activity of the upper-strand DnaB. (A and B) pBSoriC (1.6 nM) was incubated at 30°C for 9 min in the presence of IHF (16 nM), ATP-DnaA (40 nM), DnaC (100 nM), and 0, 15 or 30 nM of His-DnaB WT (WT), T86A (T), or K236A (K), followed by incubation with P1 nuclease for 5 min. The resulting DNA was analyzed by primer extension experiments, as described in Figure 8. A representative gel image from three independent experiments is shown, with annotations for each DUE subregion (A). DnaB-dependent protected region is indicated by a red bracket, and P1 nuclease-hypersensitive sites specific for WT DnaB are marked with pink arrows. Histograms of footprint profiles with 30 nM DnaB were shown (B). Similar results were obtained in the three independent experiments.

### The second DnaB loading from Right-DnaA subcomplex is required for efficient replication initiation

To further investigate the importance of the second DnaB loading for replication initiation, we conducted the form I* assay and His-DnaB pulldown assay with the Left-*oriC* plasmid (Figure 8A). The Left-DnaA subcomplex formed on Left-DOR binds to a single DnaBC complex, which should load the DnaB onto the DUE-lower strand (7,11). While stable DUE unwinding is achieved by the Left-DnaA subcomplex, both of the Left- and Right-DnaA subcomplexes are required for the full activity of the chromosome replication initiation *in vitro* and *in vivo* as well as the DnaB helicase loading (7,11). During a 9-min reaction, form I* of plasmid bearing WT *oriC* was progressively generated from form I (Figure 8B ”Input” and CD). In contrast, form I* formation of Left-*oriC* plasmid was markedly inhibited during this short time span, consistent with our previous results (7,11). Notably, in His-DnaB pulldown assay, although total recovery of the Left-*oriC* plasmid was similar to that of WT *oriC* plasmid, the form I* of the Left-*oriC* plasmid was barely recovered (Figure 8B “Pulldown” and CD).

**Figure 8.**
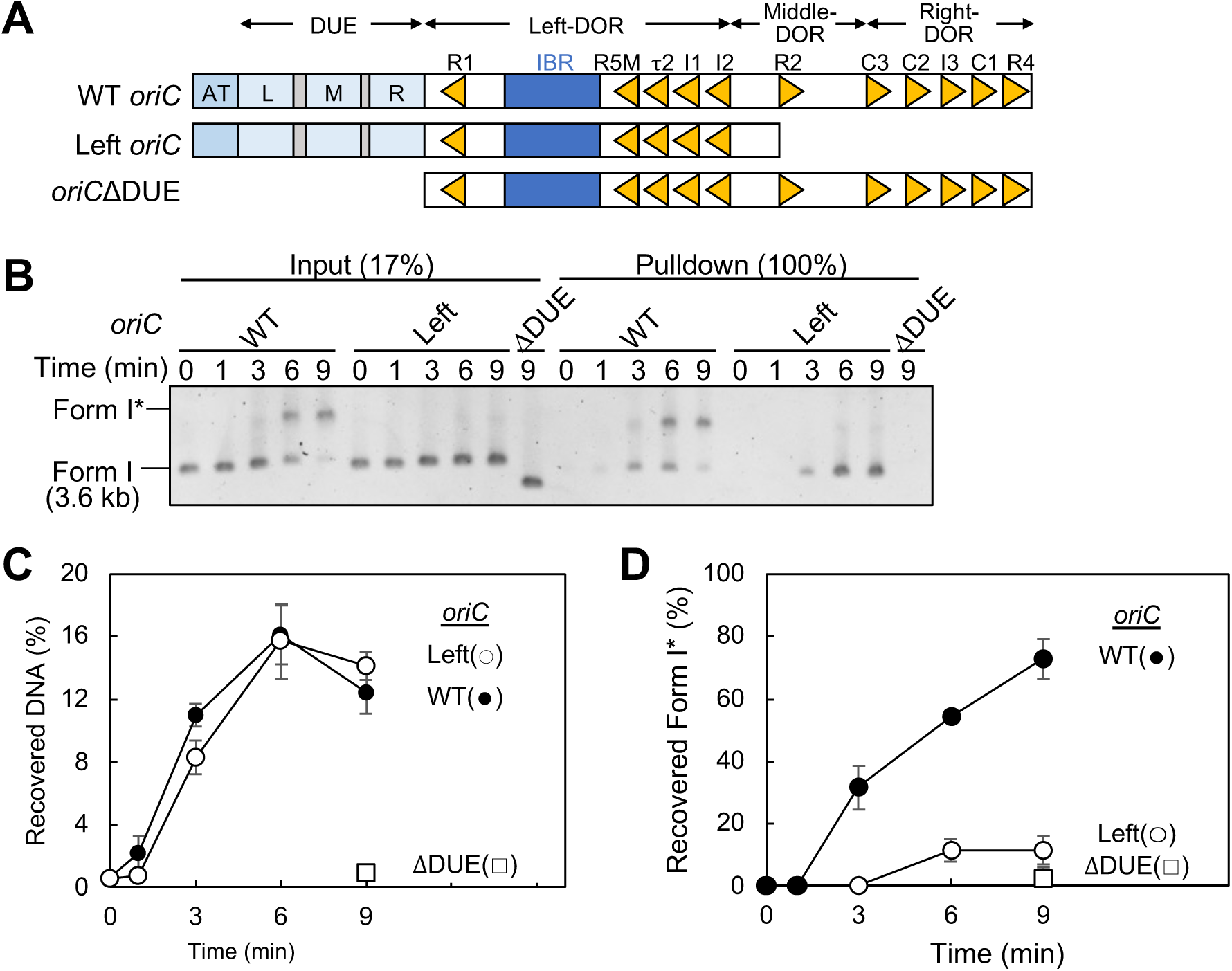
DnaB loading analysis with Left *oriC*. (A-C) His-DnaB pulldown assay. Schematic of structures of WT *oriC*, Left-*oriC* and *oriC*ΔDUE is shown (A). The supercoiled form of *oriC* plasmids (1.6 nM) bearing WT *oriC* (pBSoriC), Left-*oriC* (pBSleftoriC) or *oriC*ΔDUE (pBSoriCΔDUE) was incubated with ATP-DnaA (12 nM) with IHF (40 nM) at 30°C in the presence of His-DnaB, DnaC, gyrase, and SSB. After withdrawing a portion (25%) of the reaction mixture (Input), His-DnaB binding materials were isolated using Co2+-conjugated magnetic beads at indicated times, followed by washing with 400 mM NaCl. Input and recovered plasmid (Pulldown) were purified by phenol-chloroform extraction. Samples of 17% of the total input and 100% of the recovered plasmid were analyzed using agarose gel electrophoresis. A representative image of three independent experiments is shown in black/white-inverted mode (A). Band intensities of each lane in the gel image were analyzed by densitometric scanning. The total amounts of DNA co-recovered by His-DnaB pulldown were deduced based on the total band intensities of the “Pulldown (100%)” samples to that of the “Input (17%)” samples, and are shown as “Recovered DNA (%)” (C). The percentages of co-recovered form I* plasmid per the total co-recovered DNA were deduced from the “Pulldown” samples and are shown as “Recovered Form I* (%)” (D).

These results are consistent with our results for DnaB T86A mutant (Figure 2 and 5), and support the notion that the second DnaB loading onto the ssDUE-upper strand from the Right-DnaA subcomplex is essential for the helicase translocation creating the form I*. Considering the previous results, in the Left-*oriC* plasmid, prolonged incubation likely permits the delayed loading of the second DnaB helicase by the Left-DnaA subcomplex after the loading of the first DnaB helicase, supporting partial activity of replication initiation by Left-*oriC*. Taken together, we propose that the requirement for the second DnaB loading to support the translocation of the first DnaB helicase ensures bidirectional unwinding. This coordination between the two DnaB helicases would be of critical significance for sustaining the bidirectional replication from *oriC*.

## DISCUSSION

In this study, we investigated the specific mechanism and role for the low-affinity DnaA-DnaB interaction during replication using *in vivo* assay and *in vitro* reconstituted systems. We identified DnaB Thr86, a key residue in DnaB NTD, as being specifically required for the low-affinity interaction with DnaA. The DnaB T86A mutant was severely impaired in replication initiation activity *in vitro*, consistent with the essential role of Thr86 in cell growth and the characteristics for the DnaA His136 mutant (9). Further analyses revealed that DnaB T86A impaired its DnaB loading onto the upper strand of the unwound DUE, resulting in the loading of only a single DnaB hexamer per *oriC*. Moreover, we found that in cases of single DnaB loading, either in the presence of the DnaB T86A protein or when using the Left-*oriC* construct, translocation of the loaded DnaB was arrested. These findings reveal a novel mechanism of helicase loading and are crucial for understanding the molecular mechanism ensuring bidirectional replication from the replication origin by coordinating the loading and activation of two DnaB helicases.

Taking the present and previous reported results together, we propose a novel stepwise mechanism for DnaB loading and activation that ensures bidirectional replication initiation as follows (Figure 9). First, each of the Left- and Right-DnaA subcomplexes stably binds to a DnaBC complex through a high-affinity interaction between the DnaA domain I Phe46 and the DnaB Leu160 (7,8,10,11). The upper strand of the unwound DUE is bound by the DnaA subcomplexes (11,16). From the DnaBC complex tethered to the Left-DnaA subcomplex, the first DnaB is loaded onto the ssDUE-lower strand, which is not occupied by the DnaA subcomplex. Although the specific low-affinity interaction between the DnaA and the DnaB could contribute to this process, it is not essential. The first DnaB is initially loaded at the M region of the DUE-lower strand (Figure 6) (6,11), consistent with structural analysis indicating that the loaded DnaB covers about 20-nucleotides of ssDNA (34,54). At this step, the translocation of the loaded DnaB is arrested due to the high-affinity interaction with the multiple DnaA domain I moieties of the Left-DnaA subcomplex (Figure 9). Subsequently or simultaneously, the second DnaB is loaded onto the ssDUE-upper strand from the DnaBC complex tethered to the Right-DnaA subcomplex, in a manner dependent on the low-affinity interaction. Specifically, a DnaB protomer exposing Thr86 interacts dynamically with DnaA His136 residues exposed on the Right-DnaA subcomplex. This interaction guides the DnaBC complex precisely to the DnaA-free region of the upper strand. The strict requirement for this low-affinity interaction is likely stems from the DnaA binding to the upper strand, which restricts available space for DnaB interaction. The second DnaB is thus initially loaded at the M region of the DUE-upper strand which bears an interspace of DnaA-binding TT[A/G]T(T) sequences (Figure 6). Thus, the two DnaB are loaded on the respective ssDUE strand symmetrically around the M region, resulting in a physical collision between the two. This displaces the lower-strand DnaB from the Left-DnaA subcomplex by disrupting the high-affinity interaction with the DnaA domain I (Figure 9). Freed from the DnaA subcomplex, the DnaB helicase can translocate along the lower-strand, expanding the ssDNA region. This expanded region allows SSB binding, which in turn stimulates helicase activity and allows the upper-strand DnaB to break its own interaction with DnaA and pass through the DnaA complex (Figure 9) (46,47). As such, translocation of the first DnaB depends on the loading of the second and translocation of the second depends on that of the first. This mutual dependency ensures bidirectional unwinding around the *oriC* region, enabling bidirectional replication by sister replisomes.

**Figure 9.**
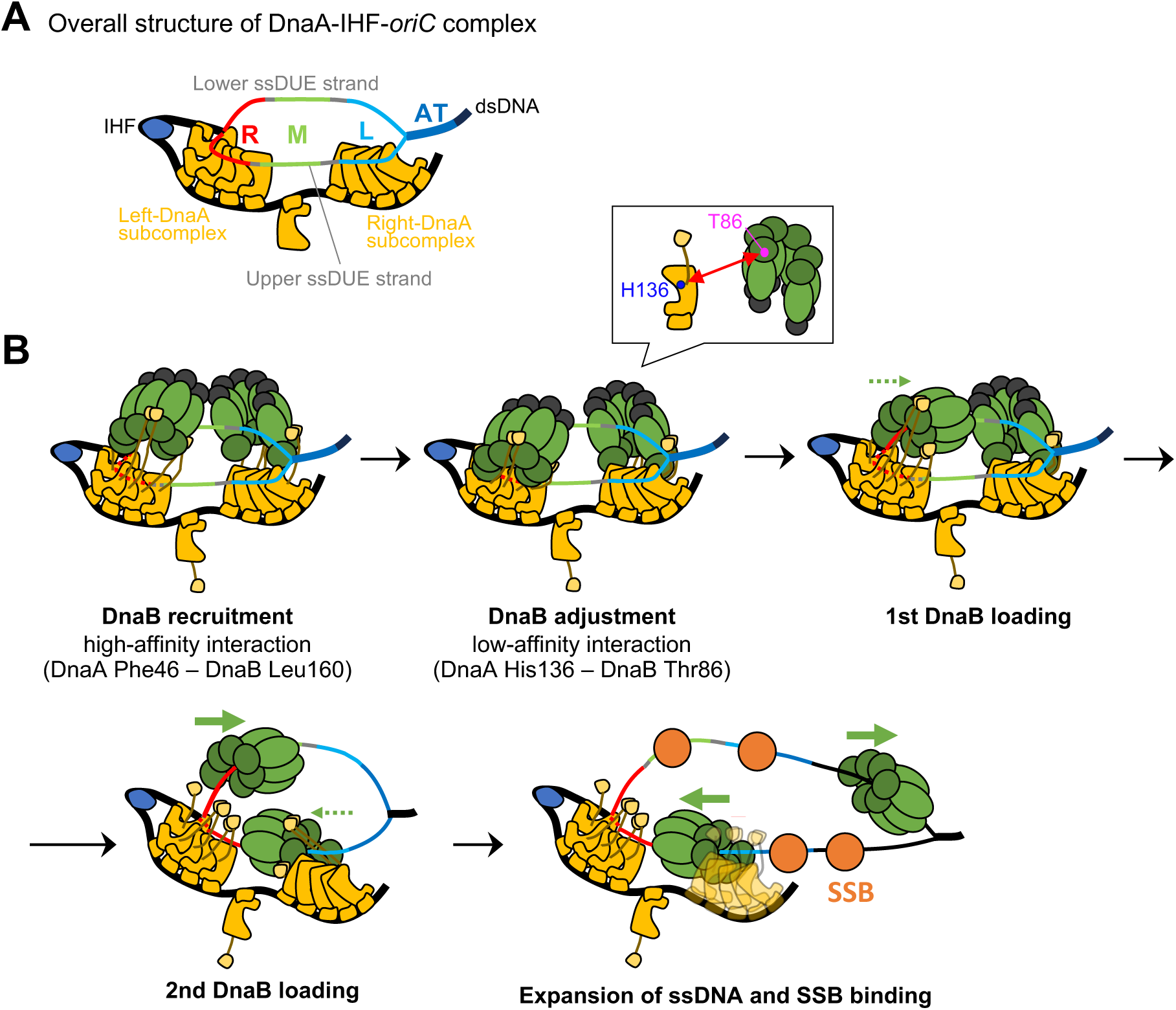
Model of strand-specific DnaB loading mechanism during bidirectional replication initiation. (A) A schematic representation of the expanded open complex. Proteins are indicated as those in Figure 1D. The L/M/R-DUE and AT-cluster regions are indicated by different colours. (B) A schematic representation of the coordinated mechanisms in DnaB loading and translocation. The *oriC*-IHF-DnaA complex unwinds the DUE forming the expanded open complex, and each DnaA subcomplex engages a DnaBC complex via the high-affinity interaction (*upper left panel*). Subsequently, the low-affinity interactions between the DnaA H136 and the DnaB T86 residue (*insert*) take place, while the high-affinity interaction is maintained (*upper middle panel*). The Left-DnaA subcomplex first loads a DnaBC complex onto the M region of ssDUE-lower strand with the moderate but non-essential support from the low-affinity interaction (*upper right panel*). Meanwhile, the Right-DnaA subcomplex loads the second DnaBC complex onto the upper strand of the unwound M-DUE region in a manner dependent of the low-affinity interaction (*lower left panel*). Even after dissociation of DnaC from the loaded DnaB, physical collision occurs between the first and the second DnaB hexamers. This collision leads to dissociation of the first DnaB from the link with the DnaA subcomplex, enabling its translocation. The expanded ssDUE region generated by the first DnaB recruits SSB molecules, which stimulates DnaB helicase activity, thereby promoting the translocation of the second DnaB. These coordinated events ensure bidirectional progression of DnaB helicases (*lower right panel*).

In the Left-DnaA subcomplex, DnaA domain III H/B-motifs stably bind the DnaA-binding motifs, TTATT and TTGT sequences, in the upper-strand DUE-R/M region (Figure 1) (11,15,20). In the above hypothesis, the first DnaB is loaded onto the ssDUE-lower strand, which is free from DnaA binding. Although the specific low-affinity interaction between DnaA and DnaB should help guide the DnaBC complex, the DnaB tethered to the Left-DnaA subcomplex would be preferentially loaded onto the DnaA-free ssDUE strand in the close vicinity even without the low-affinity interaction.

In the Right-DnaA subcomplex, DnaA domain III H/B-motifs should bind the DnaA-binding motifs, TTATT sequence, in the upper-strand DUE-L region (Figure 1) (11). A previous study suggest that this interaction is relatively unstable but is effective to expand the unwound region, optimizing the loading processes of the two DnaB helicases onto ssDUE region (11). In the above hypothesis, the second DnaB tethered to the Right-DnaA subcomplex is loaded onto the upper ssDUE strand which binds to the Right-DnaA subcomplex. In this process, the low-affinity interaction between the DnaA domain III His136 and the DnaB Thr86 is required, which likely guides the DnaB to the interspace of the DnaA-binding sequences within the M region of ssDUE. Given that binding of ssDUE to the Right- DnaA subcomplex is relatively unstable (11), L-ssDUE region should be frequently dissociated from the DnaA complex in an equilibrium manner, which should stimulate the second DnaBC complex to capture the DnaA-free ssDUE to load the DnaB onto the ssDNA. The low-affinity interaction of the DnaB with DnaA would be required to keep the DnaBC complex in the close vicinity to the ssDUE interacting with the Left-DnaA subcomplex, which ensures timely interaction of the DnaBC complex with ssDUE dissociated from the Right-DnaA subcomplex. Alternatively, the low-affinity interaction could change structure of the Right-DnaA subcomplex, stimulating the releasing of ssDUE and its interaction with the second DnaBC complex.

Based on our results and previous structural analyses, we propose that a specific protomer within the DnaB hexamer is essential for the low-affinity DnaA-DnaB interaction. Binding of DnaC to DnaB induces a conformational change that converts DnaB into an open helical state (34,52). At this stage, a specific DnaB protomer located at the ring-opening site adopts a protruding conformation of its NTD, prominently exposing Thr86 on the surface (Supplementary Figure S2). Given that the DnaA His136 residues are exposed and regularly aligned on the DnaA subcomplexes (9,19), it is plausible that the protruded Thr86-containing NTD of this specific DnaB protomer dynamically interacts with the DnaA His136 in the complex (9,19). This hypothesis is supported by DnaC-dependent recovery of DnaB observed in our bio-*oriC* pulldown assay using the DnaA F46A mutant (Figure 4). According to this model, the specific interaction between the ring-opening site of DnaB hexamer and the DnaA subcomplex would play a critical role in regulating the orientation and positioning of DnaB in an optimal conformational state for engaging ssDUE.

Consistent with the above proposed mechanism (Figure 9), structural model of the loading intermediate DnaA-DnaBC complex with the low-affinity interaction can be constructed using the known three-dimensional structure of the DnaBC complex and the structural model of the DnaA-IHF-*oriC* complex derived from the molecular dynamics simulations (Supplementary Figure S3) (19,34). Based on the aforementioned considerations and the specific role for R5M and R4 DnaA boxes in replication initiation and helicase loading (11,16,20), we assume the interaction between the His136 residues of the R5M- and R4-bound DnaA protomers and the Thr86 residues of the DnaB protomer adjacent to the ring-opening site (Supplementary Figure S3). Interestingly, this interaction potentially positions the ring-opening site of DnaB in the close vicinity to the ssDUE, supporting that the results from biochemical analysis and the suggested model for the bidirectional loading of DnaB are consistent with the nucleoprotein complex structure and such a loading intermediate has a reasonable configuration to promote DnaB loading onto the ssDUE (Figure 9).

The result of the complementation assay suggests that the methyl group is crucial for the function of the Thr86 residue (Table 2). Thr86 is highly conserved among bacterial DnaB, and its surrounding region also contains conserved valine, leucine, and isoleucine residues, all of which possess methyl groups (Figure 1C). In DnaA orthologs of various bacterial species, the amino acids at the position corresponding to *E. coli* DnaA His136 are conserved in those with an aromatic moiety such as phenylalanine and tyrosine as well as histidine (9). Therefore, a low-affinity interaction between the aromatic amino acids in DnaA domain III and the 4-methyl-containing amino acids in the DnaB NTD may be widely conserved among bacteria.

Finally we suggest that the stepwise DnaB loading mechanism including the helicase- ring opening site-specific interaction with DnaA represents a primordial mechanism for replicative helicase loading at the replication origin. In eukaryotes, the ORC-Cdc6 complex at the origin loads the Cdt1-MCM helicase complex onto double-stranded DNA (termed as OCCM) (12). Recent studies have reported that the Cdt1-MCM helicase complex is loaded in a stepwise and dynamic manner through multiple intermediates referred to as “semi-attached OCCM” and “pre-insertion OCCM” (13,14). Cross-linking/mass spectrometry analysis of the yeast OCCM has suggested that Mcm2 and Mcm5 subunits, located at the helicase-ring opening site, interact with Orc subunits during the process of construction of OCCM (13). Taking the site-specific ORC-MCM interaction occurring at the MCM-ring opening site into consideration, the present study of *E. coli* DnaB helicase loading mechanisms may represent a common principle in the fundamental helicase loading mechanism onto the origin DNA.

## SUPPLEMENTARY DATA

Supplementary Data are available.

## DATA AVAILABILITY

All data are included in the article and in Supplementary Material.

