## Supplementary for "Dynamic DnaA-DnaB interactions at *oriC* coordinate the loading and coupled translocation of two DnaB helicases for bidirectional replication"

**Supplementary Table S1. Oligonucleotide used in this study**

| Name | Sequence | note |
|---|---|---|
| R1UP | TTTTTTTTTTTTTTTTTTTTTTTTTTTTTTTTTTTGCCCTGTGGAT AACAAGGATCCGGCTTTTAAGATCAACAACCTGGAAA | Used for helicase activity assay |
| R1LO | TTTCCAGGTTGTTGATCTTAAAAGCCGGATCCTTGTTAT CCACAGGGCATTTTTTTTTTTTTTTTTTTTTTTTTTTTTTTTTTT | |
| R1LOcomp | TTTCCAGGTTGTTGATCTTAAAAGCCGGATCCTTGTTAT CCACAGGGCA | |
| ori1 | ATCGCACTGCCCTGTGG | Used for construction of bio-DOR fragment |
| ori-2 | CAAATAAGTATACAGATCGTGCG | |
| KWSmaloriCFwd | CCCGGGCCGTGGATTCTAC | Used for primer extension |
| Dr2-2 | CCCCTCATTCTGATCCCAGC | |

**Supplementary Table S2. *oriC* plasmids used in this study**

| Name | Relevant structure | Reference |
|---|---|---|
| pBSoriC | pBluescript II bearing oriC | Sakiyama, et al., 2022 |
| pBSoriCΔDUE | pBSoriCΔDUE | Sakiyama, et al., 2022 |
| pBSleftoriC | pBSoriC Δ(R4-R2) | Sakiyama, et al., 2022 |

**Supplementary Figures**

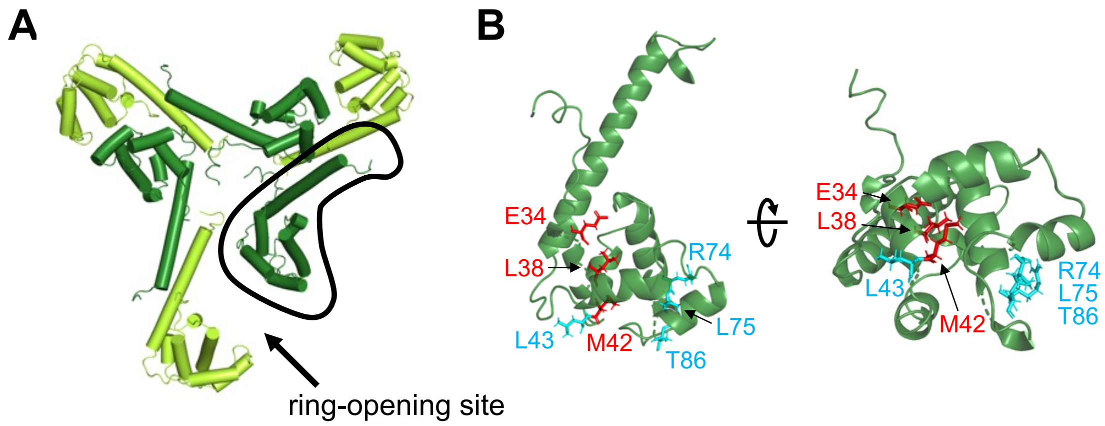


**Supplementary Figure S1 Structure of the DnaB N-terminal domain**

(A) Top view of the DnaB NTD within the cryo-EM structure of DnaB-DnaC complex (Arias-Palomo et al., 2019) (PDB: 6QEL). Each protomer is alternately shown in green or light green colors.

(B) Close up of the DnaB NTD outlined in (A). Among the essential residues shown in Figure 1B or Table 1, those likely exposed on the protein surface are indicated in light blue, while those facing inward are shown in red.

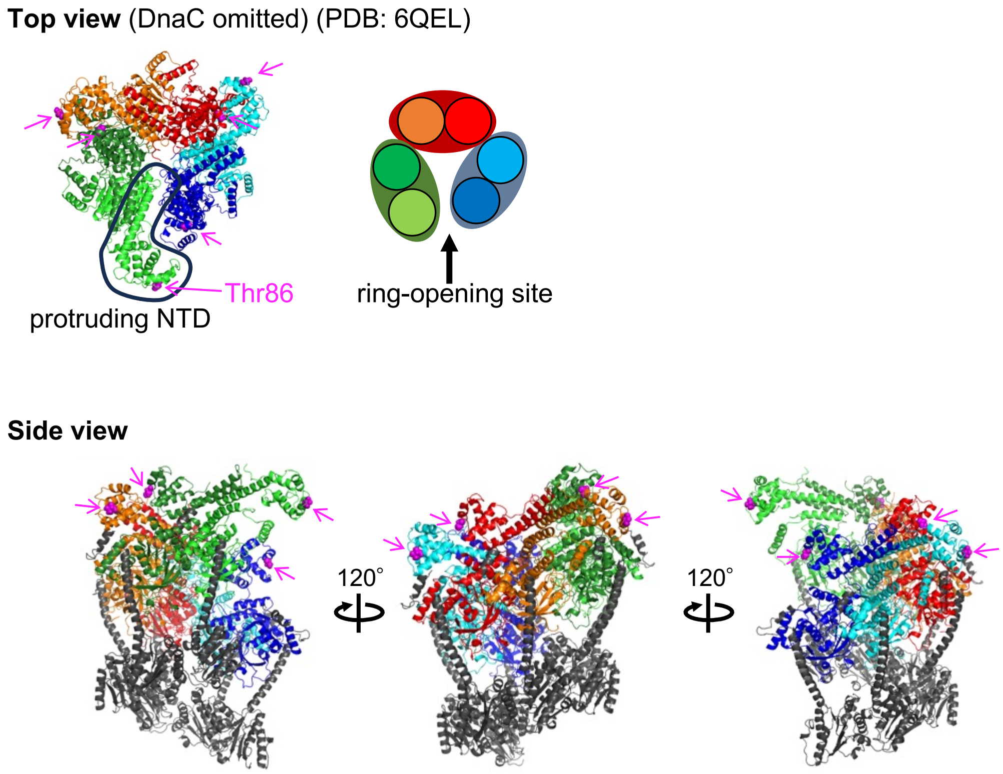


**Supplementary Figure S2  Structure of each DnaB subunit within the DnaB–DnaC complex**

Top and side views of the DnaB-DnaC complex (Arias-Palomo et al., 2019) (PDB: 6QEL). Each DnaB protomer is indicated in different colors as illustrated. DnaC is indicated in dark gray. Thr86 residues in each DnaB protomer are represented in magenta. A specific protomer at the ring-opening site with distinctive protruding conformation is outlined (see discussion).

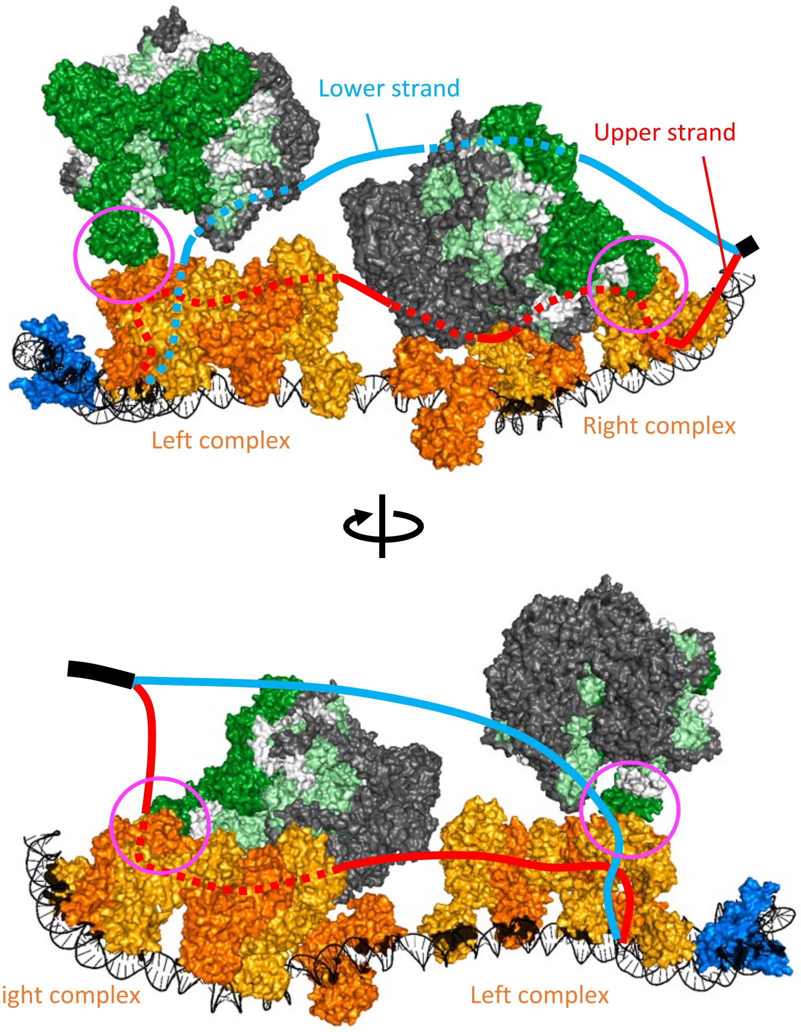


**Supplementary Figure S3 Structural model of the loading intermediate DnaA-DmaBC complex with the low-affinity interaction**

The structural model of low-affinity DnaA-DnaB interaction for DnaB loading on *oriC* is manually reconstituted using the cryo-EM structure of DnaB-DnaC complex (Arias-Palomo et al., 2019) (PDB: 6QEL) and the structural model of the DnaA-IHF-*oriC* complex derived from molecular dynamics simulations (Shimizu et al., 2016) with PyMOL. In the molecular dynamics simulations, the whole structure of DOR and IHF in addition to DnaA domains III-IV are adopted. DnaB and DnaC are colored as in Figure 9, except that DnaB LH domain is indicated in white. The lower and upper strands of the region spanning DUE to the AT-cluster are indicated by light blue and red lines, respectively. IHF is shown in blue colour. DnaA protomers are alternately shown in yellow and dark yellow colours. The interaction regions between DnaB Thr86 and His136 of DnaA bound to R5M and R4 boxes are indicated by magenta circles.
